# Mapping Spatially Distributed Material Properties in Finite Element Models of Plant Tissue Using Computed Tomography

**DOI:** 10.1101/2020.02.27.968651

**Authors:** Christopher J Stubbs, Ryan Larson, Douglas D Cook

## Abstract

Plant tissues are often heterogeneous. To accurately investigate these tissues, we will need methods to spatially map these tissue stiffness values onto finite element models. The purpose of this research is to develop a method for using specimen-specific computed tomography data to inform the spatial mapping of Young’s modulus values on finite element models. The spatial mapping of Young’s modulus was calculated and then used to predict the response of specimen tests. Results indicated that this method can be used to obtain spatial distributions of material properties, thus enabling finite element models that account for material heterogeneity.

## Introduction

Biological materials typically exhibit spatial variation of mechanical properties (Wimmer et al. 1997; Gourion‐Arsiquaud et al. 2009; Manjubala et al. 2009). In addition, mechanical properties of the material often covary with other physical measurements of the material. When these covariance patterns are known, estimates of one property can be obtained from measurements of a different property. This technique has been used in human and animal tissue biomechanics, such as estimating the Young’s modulus of a material based on its second harmonic autofluorescence (Liu et al. 2019) or nanoindentation response (Wimmer et al. 1997; Zysset et al. 1999; Cuy et al. 2002).

Cellular materials such as plant tissues often show a linear relationship between the mechanical properties of the material and the density of the material (Gibson 2005). And the intensity of x-ray computed tomography (CT) data generally correlates with material density [REFS needed]. This suggests that CT intensity could be used to infer the distribution of mechanical tissue properties. Indeed, this approach has been successfully employed to model the heterogeneity of bone (Helgason et al. 2008), where a linear relationship between CT intensity and mechanical properties of human bone provides reasonable results (Ciarelli et al., 1991; McBroom et al., 1985; Rho et al., 1995). These relationships have been used to create finite element models using a variety of CT-informed material mapping algorithms (Taddei et al., 2004). These methods and relationships developed in the field of human and animal biomechanics can be very valuable to the field of plant biomechanics. As a less developed field, plant biomechanics can accelerate the research field by leveraging these methods and successfully modifying them to be applicable in plants.

The first purpose of this study was to assess the feasibility of using CT data to infer the spatial distribution of the transverse Young’s modulus within cross-section of maize stem specimens. The second purpose was to assess the validity of such a mapping relationship through secondary tests to validate the predictive utility of the mapping relationship.

## Materials and Methods

### Compression testing

Commercial hybrid maize stems were used as test specimens for this study. Stems were sectioned into specimens that were approximately 10mm in length (see Figure 1). Each specimen was tested in transverse compression, loaded through the minor diameter of the specimen cross-section. Next, the pith was removed from each specimen, and they were tested in the same loading configuration. Linear force-displacement curves were extracted from each test. Further details of the experimental design, including test equipment, sample preparation, etc. were documented in a previous study (Stubbs et al. 2019).

**Fig. 1.**
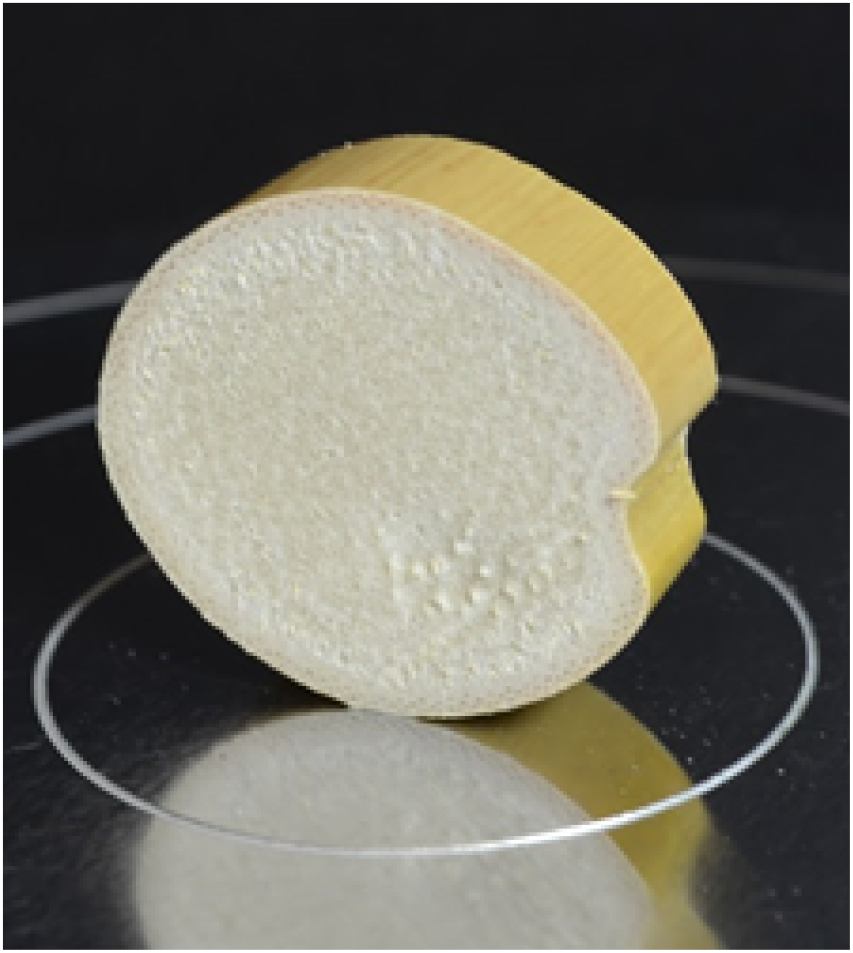
A typical compression test specimen (Stubbs, 2019).

### Finite Element Model Development

A finite element model was developed to simulate the testing in Abaqus/CAE 2016. The geometry for the model was created using the inner and outer boundaries derived from the specimen-specific CT scans, as described in a previous study from our research group (Al-Zube et al. 2017). The model was designed as a two-dimensional plane-stress analysis, with transversely isotropic material properties. The model was meshed using 4-noded bilinear plane stress quadrilateral elements (Hibbitt et al. 2016; Simulia 2016). For the analyses that simulated the hollow specimen test configuration, the pith elements were removed.

The finite element models were solved using a non-linear, full Newton direct solver in Abaqus/Standard 2016. The contact between the platens and the specimen was represented by a general contact interaction with a finite-sliding formulation and a hard penalty pressure-overclosure relationship. Tangential friction was used to aid in convergence with a negligibly small coefficient of friction. Further details of the modeling approach were documented in a previous study (Stubbs et al. 2019).

### CT mapping of material properties

A mapping equation was required to relate CT data to the finite-element model. The CT scan data consisted of a 3-dimensional array of x-coordinate, y-coordinate, and an 8-bit grayscale value representing the CT intensity of the pixel. A linear relationship between CT intensity and modulus was assumed. The Young’s modulus values were mapped to the FEM using a USDFLD FORTRAN subroutine in Abaqus/Standard 2016. Figure 2 depicts the original CT scan data and the inferred Young’s modulus data. The Poisson’s ratio was set to a value of 0.25 for all analyses, as Poisson’s ratio was previously found to have a negligible effect on this type of analysis (Stubbs et al. 2019).

**Fig. 2.**
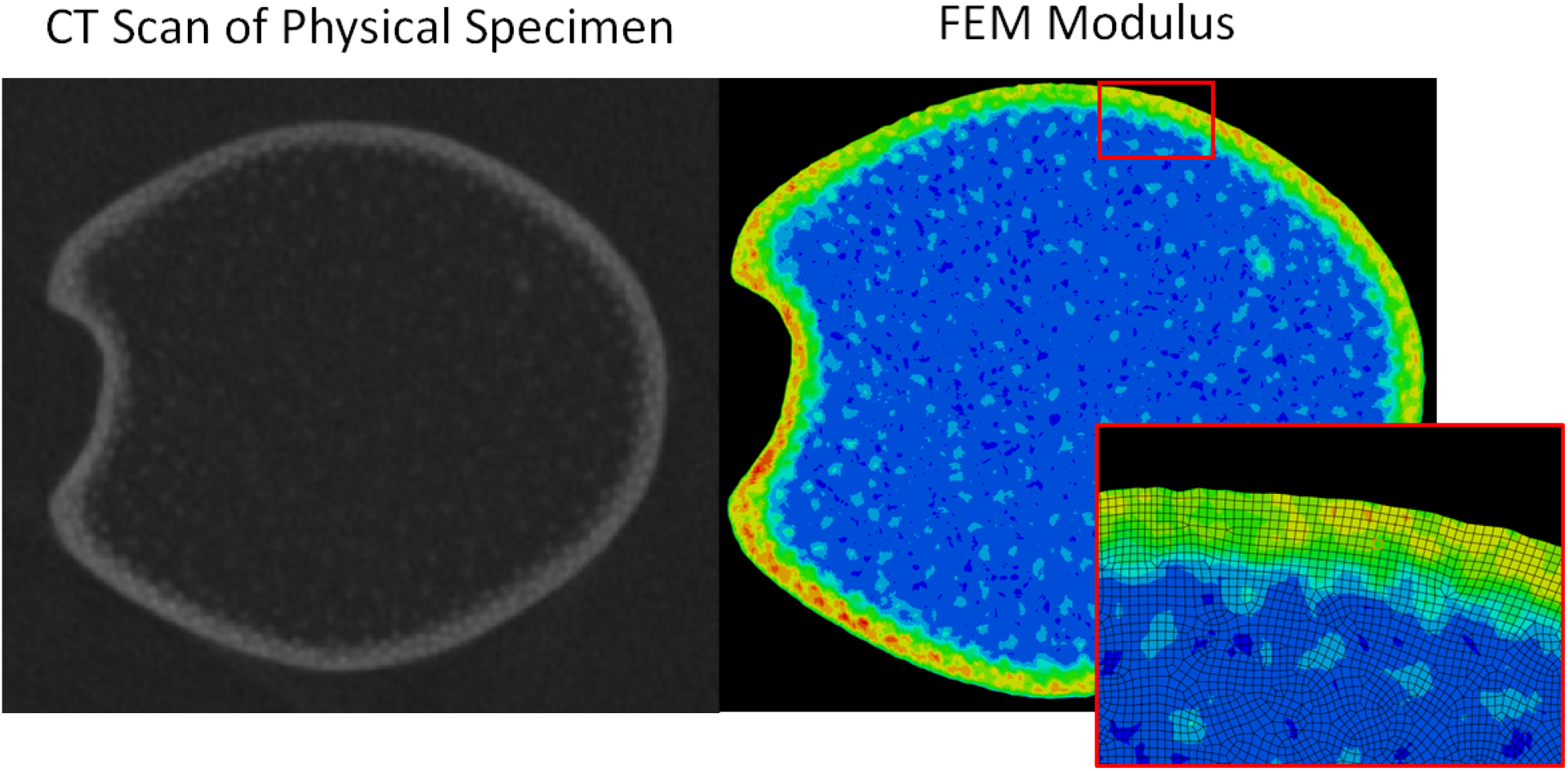
Left: A representative CT scan of a single specimen. Right: the corresponding finite element model, mesh, and (in color) the inferred CT intensity values of each element.

A mapping equation that relates CT intensity with Young’s modulus was defined by a linear function with a slope *S*. In addition, there exists a lower bound CT intensity value (*CT_min*) at which the Young’s modulus is 0. Naturally, to prevent non-physical behavior, all CT intensity values less than this value would also correspond to a Young’s modulus of 0. As there exist two unknowns in this function (*S*, *CT_min*), we needed to create a system of two analyses to simultaneously solve to determine these values. These two analyses correspond to the two tests performed on each specimen. In short, determining the CT mapping function was done by solving for the coefficients of a linear mapping function that is capable of accurately reproducing the structural response of both physical tests of a given specimen.

To solve for the coefficients (*S* and *CT_min*), the following procedure was used: (1) three initial estimates were made of *CT_min* values for each of the two specimen tests, (2) the three corresponding *S* values were iteratively solved for in each of the two specimen tests, (3) quadratic approximations were found for the two *S* vs. *CT_min* curves, (4) an estimate was made of the *CT_min* that would result in both analyses having the same *S* values (the point at which the two curves intersect), (5) the two analysis were both iterated upon at this newly predicted *CT_min* value, and the resulting *S* values were compared. If the two *S* values were found to be within 1% of each other, the mapping equation was considered to be valid. Otherwise, steps 2 through 4 were repeated. Graphical representations of this process are shown in Figure 3.

**Fig. 3.**
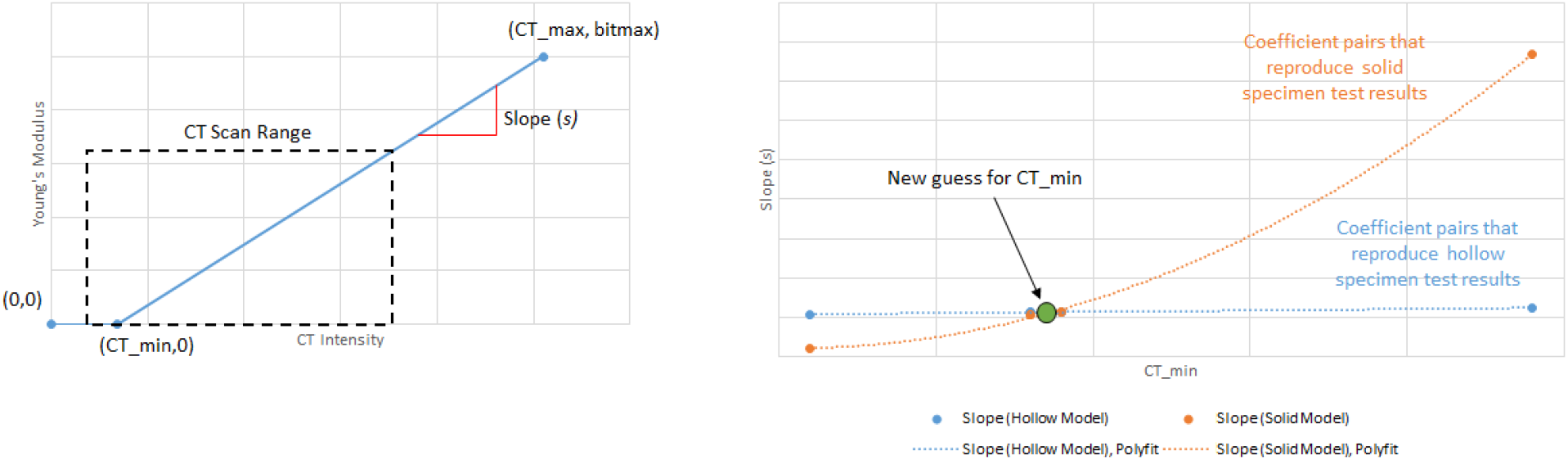
Left: The mapping function relating CT intensity to Young’s modulus, including the two unknown values CT_min and slope, and the range of CT intensity values found in a typical CT scan of a maize stem specimen. Right: an example of the valid CT_min and slope pairs for a specimen (right). Quadratic approximations for the two curves were found and iterated to find the intersection of the curves, i.e. the CT_min and s pair that resulted in a CT intensity mapping equation that was valid for both test configurations.

### Validation

Mapping relationships were validated using a completely independent data set. This was accomplished by testing each specimen in a different orientation: with the compressive load applied along the major axis of the cross-section, as shown in Figure 4. As the validation testing was performed with the pith intact, these tests were performed prior to the pith removal described in the previous sections. A new finite-element model was built to simulate this loading configuration, and the CT-mapping equation was applied to the model. The predicted force-displacement structural response of the model was then compared to the force-displacement slope from the physical test.

**Fig. 4.**
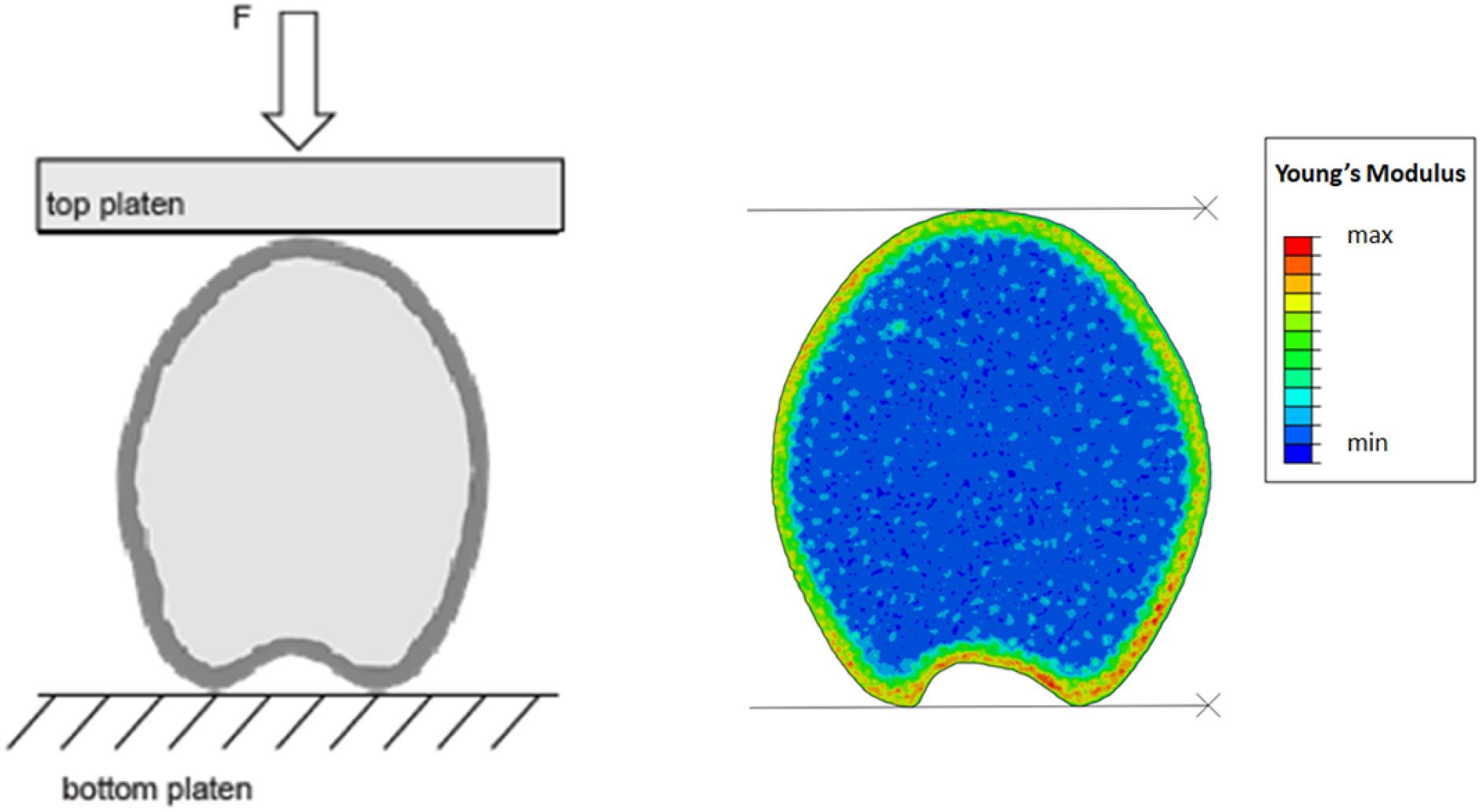
The validation testing configuration (left) (Stubbs et al. 2019); the validation FEM (right).

## Results

### Mapping Coefficients

Mapping coefficients were successfully determined using the approach described above. The coefficients of each sample’s mapping functions are plotted in Figure 5, with *CT_min* values along the horizontal axis and slope values along the vertical axis. In this chart, specimens from the same maize stem are plotted using the same type of symbol (e.g. circle, x).

**Fig. 5.**
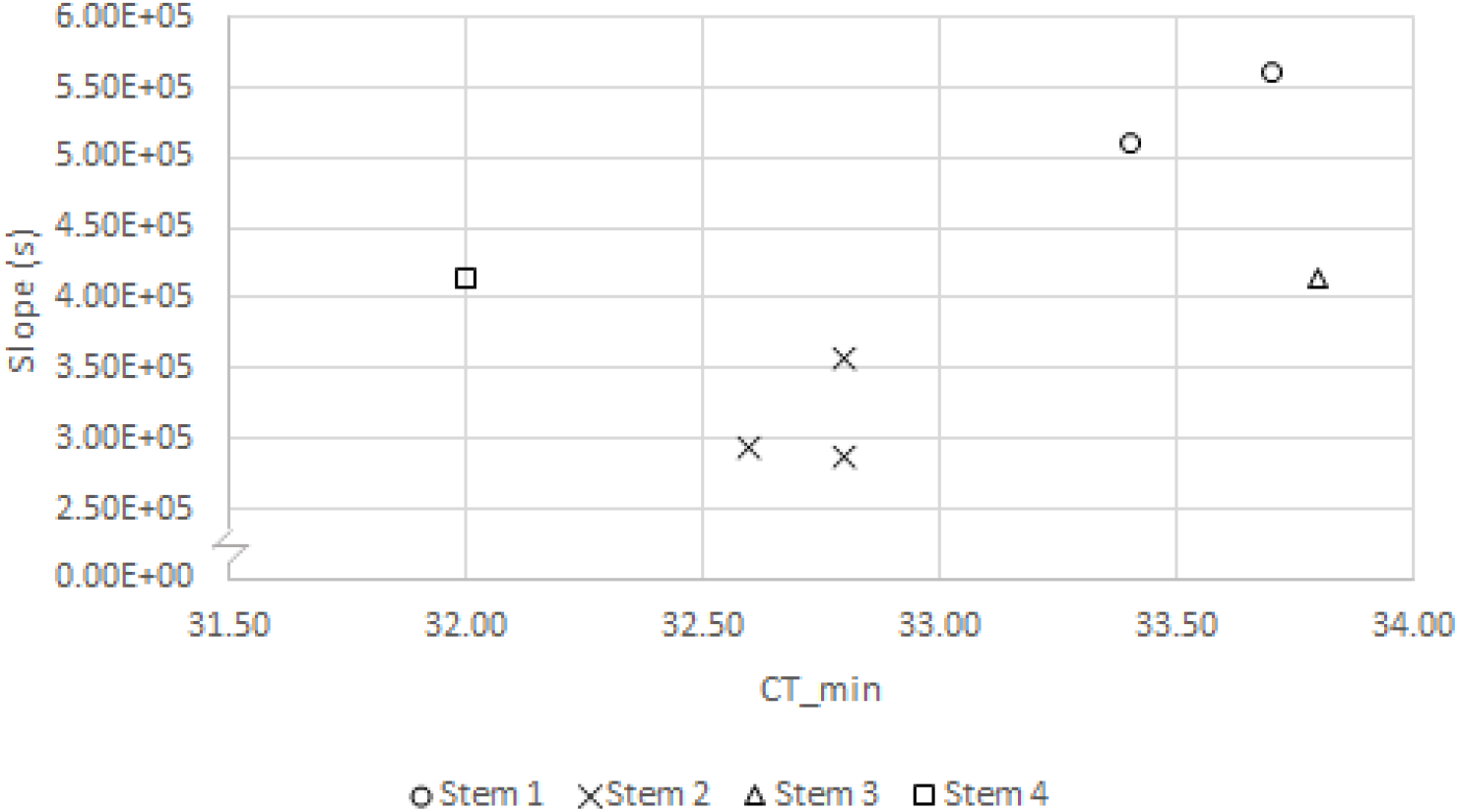
The CT_min and slope values of the mapping functions of the specimens.

While the *CT_min* coefficient values were quite consistent (coefficient of variation of 1.6%), there was a much higher variation in the slope coefficient (coefficient of variation of 25.4%). This was somewhat unexpected, as all of the specimens were imaged within a single CT scan, suggesting that both solution parameters would have had similar coefficients of variation. This issue is addressed further in the discussion section.

### Independent Validation

The method for obtaining CT-mapping coefficients ensured that, for each specimen, the modeled vs. measured structural response of solid and hollow specimens were both within 1%. This itself could be considered as a type of validation since a single mapping relationship could be used to accurately predict the response of the specimen in two different configurations (solid and hollow). However, we also tested the validity of the mapping relationships and their predictive utility using secondary tests. This was done by creating models of the validation tests in which the load was applied along the major axis of the cross-section (Figure 4). The validation models predicted the structural response of the specimen tests with an average error of 7.85% and a median error of 2.01%. The comparison between measured and predicted structural responses of each validation model is plotted in Figure 6. The ability of the models to accurately predict the force-displacement response of the specimen in a different test configuration was considered to be positive validation of the CT-mapping method.

**Fig. 6.**
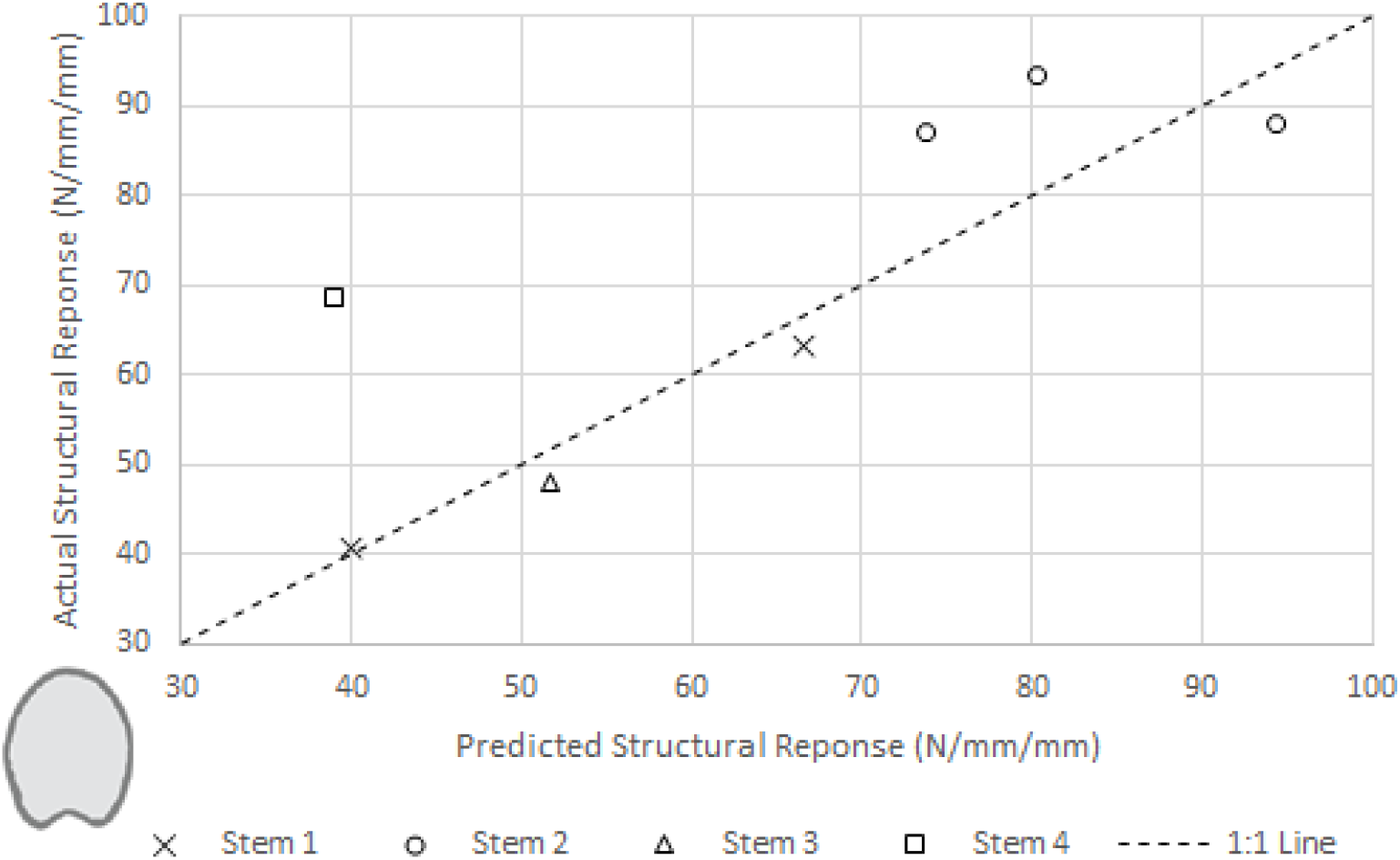
The predicted vs. actual structural response of the validation models, with 1:1 line shown for reference.

**Fig. 7.**
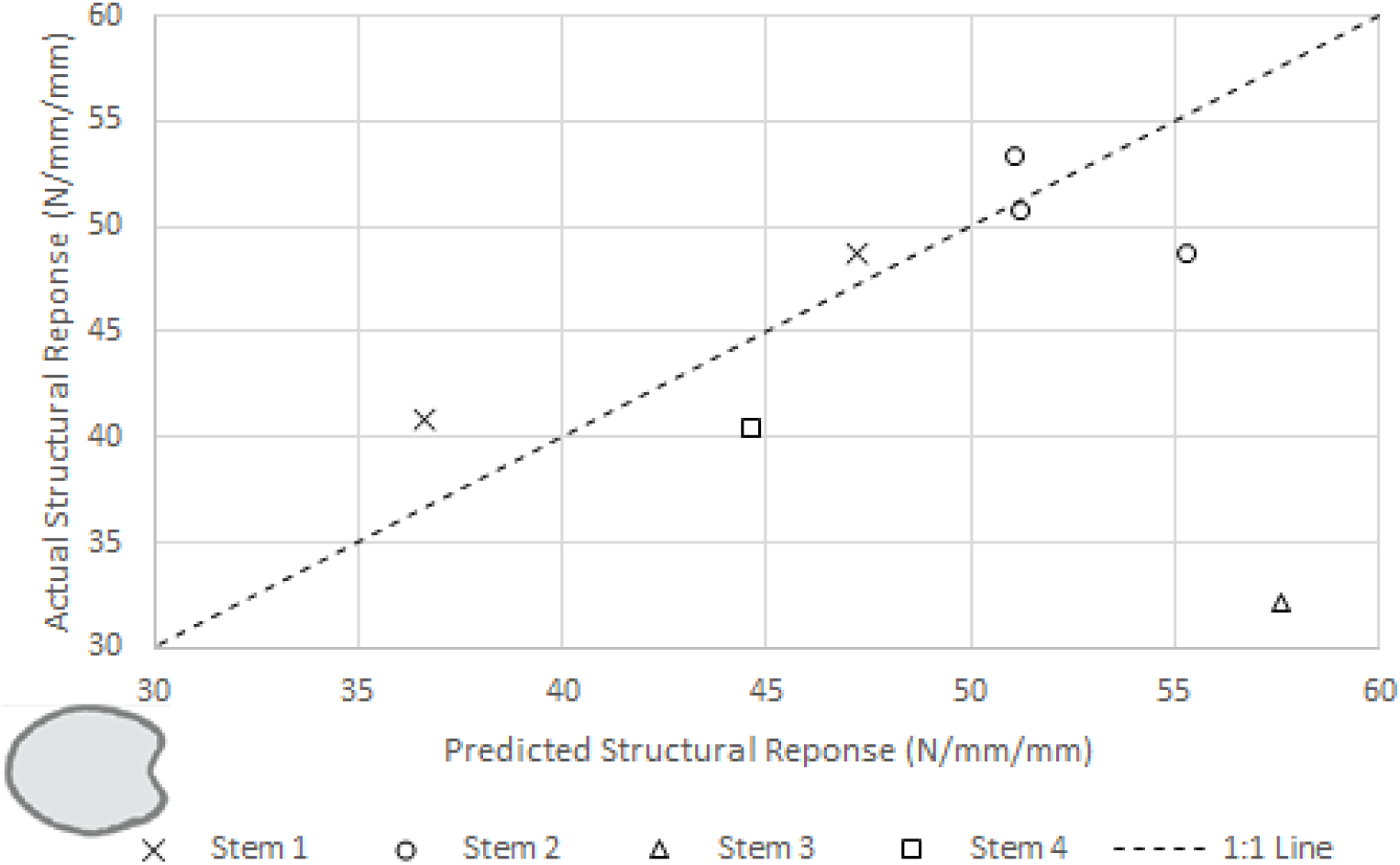
The predicted vs. actual structural response of the models when using the averaged mapping function values, with 1:1 line shown for reference.

### The Utility of Average Mapping Coefficients

Using the CT-mapping coefficient data from Figure 5, average *CT_min* and slope values were calculated. The average values were *CT_min* = 33.0 and *S* = 4.05E5. The models were then re-analyzed in their original configuration (load applied along the minor axis of cross-section) to determine the error levels introduced by the use of average CT-mapping coefficients. The average CT-mapping relationship predicted the structural response of the specimen tests with an average error of 12.3% and a median error of 0.84%. A comparison of the predicted structural response using averaged mapping vs. the actual structural response of each specimen is plotted in Figure 6. These data are encouraging since the use of a single mapping function produces reasonable levels of error across most specimens. This suggests that a single mapping function can potentially be applied across an entire scanned stalk.

## Discussion

### Variation in Coefficients and Predictive Accuracy

The method used in this study produces a single pair of CT-mapping coefficients from each specimen. This approach was used because (a) we anticipated that mapping coefficients would be relatively similar across specimens, and (b) because it provides a very simple and straightforward solution process that minimizes the number of finite-element solutions required to obtain the mapping coefficients. The resulting coefficients provide a unique CT-mapping relationship for each specimen. This specimen-specific mapping relationship is highly accurate at predicting the solid and hollow responses of each individual test specimen, and has a relatively low level of error (median error of 2%) when each specimen is tested in a different configuration. Even though all specimens were scanned together in the same scan, the slope coefficient values exhibited a wider level of variation than expected. But on the other hand, when average coefficient values were used, most models exhibited low levels of error (median error of 0.84%).

This study was focused on a first attempt to obtain CT-mapping relationships for plant materials, and the results are encouraging. But the results are not without drawbacks and limitations. We have identified several possible causes that might explain the range in slope coefficients that was observed in Figure 4. These possible causes include (a) repeatability errors in the experimental test data, (b) inconsistencies in CT-scan data, (c) longitudinal variation in the material properties of actual test specimens that are not captured in a two-dimensional model, (d) the solution approach used in this study provided specimen-specific coefficient pairs instead of a single pair of coefficients that was simultaneously optimized across the entire set of specimens, or (e) a combination of these factors. Further analyses will need to be performed to determine the influence of each of these potential causes. Such future research should indicate whether the variation observed in this study is due to inherent limitations in measurement techniques, or if the variation could perhaps be reduced by refinement of the measurement method. In addition, as this data set represents a limited number of samples taken from only four stems, further investigation will be required to more carefully examine how the mapping function changes between stems and along the entire length of stems.

### Uses of CT-mapping Relationships

First, this method provides a new way to create models of plant behavior that account for the variation in material stiffnesses. This should enable a deeper exploration of the stress states within a plant tissues than is possible when using simple, homogenous material modeling (Hammond et al. 2018). These results were obtained in Abaqus 2017. But readers should be aware that scan data can now be directly imported into the model as analytical fields in Abaqus/CAE 2019. This negates the need for a custom user subroutine, and drastically reduces the computational cost of this type of simulations. Thus, such simulations are now much faster and more user-friendly.

### Limitations

This study was conducted to determine if CT intensity data could be used to infer the transverse Young’s modulus on a per-specimen basis. The results suggest that this approach is possible, and provides reasonably accurate results. However, the method is not without limitations. There are several limitations of this study and possible improvements. For example, CT-scanning is typically only feasible using dried specimens because water tends to absorb much more x-ray radiation than the dry matter of a stem. This is less of a limitation in the study of maize stems since we are often interested in the failure of maize stems just before harvest when they are relatively dry. Second, in this study we assumed a linear relationship between CT intensity and material stiffness. While the density-stiffness relationship of cellular materials is well-established (Gibson 2005), x-ray absorption is a relatively complex phenomenon which is likely not exactly linear in nature. Third, the validation approach used in this study was dependent upon overall specimen outcomes. The close agreement between predicted and measured response is certainly encouraging, and is likely correct. However, the mapping relationship itself was not tested *directly*. The authors have previously considered the use of indentation techniques to assess the local *longitudinal* stiffness of maize tissues, but at the microscale, maize tissue is highly inhomogeneous. This makes such approaches very problematic. As more sophisticated material assessment techniques become available, it may be possible to more directly assess the accuracy of the mapping relationships obtained in this study. Fourth, only a single transverse CT scan layer was used for each specimen and the finite-element model was two-dimensional. This approach ignores any changes in the transverse cross-section along the length of the specimen. Further improvements to the method could be obtained through taking into account changes in the cross-section and the spatial distribution of tissue stiffness along the length of the specimens.

## Conclusion

A method was developed for inferring the spatial distribution of the transverse Young’s modulus within cross-section of maize stem specimens form CT scan data. The method essentially eliminates error on a per-specimen basis under initial loading conditions, and when these same specimens were tested in secondary configurations, the CT-informed material mapping was able to predict specimen behavior with an average error of 7.85%. When using a single average mapping relationship, the average error across all specimens was 12.3%. These results indicate that CT scan data can be used to infer the spatial distribution of transverse Young’s modulus within plant specimens, an approach which will enable more detailed studies that consider material heterogeneity.

## Geolocation Information

This research was performed at New York University Abu Dhabi, (Abu Dhabi, United Arab Emirates), New York University Tandon (New York, NY), Brigham Young University (Provo, UT), and University of Idaho (Moscow, ID).

## Acknowledgements

We thank Bayer AG (formerly Monsanto Company) for providing the maize stem samples used in this study. This work was funded in part by the National Science Foundation (Award # 1400973).

